# Improved estimation of molecular evolution coupling stochastic simulations and deep learning

**DOI:** 10.1101/2023.09.08.556821

**Authors:** Daniele Silvestro, Thibault Latrille, Nicolas Salamin

**Affiliations:** Department of Biology, University of Fribourg and Swiss Institute of Bioinformatics, 1700 Fribourg, Switzerland; Gothenburg Global Biodiversity Centre, Department of Biological and Environmental Sciences, University of Gothenburg, 40530 Gothenburg, Sweden; Department of Computational Biology, University of Lausanne, 1015 Lausanne, Switzerland

## Abstract

Models have always been central to inferring molecular evolution and to reconstructing phylogenetic trees. Their use typically involves the development of a mechanistic framework reflecting our understanding of the underlying biological processes, such as nucleotide substitutions, and the estimation of model parameters by maximum likelihood or Bayesian inference. However, deriving and optimizing the likelihood of the data is not always possible under complex evolutionary scenarios or tractable for large datasets, often leading to unrealistic simplifying assumptions in the fitted models. To overcome this issue, we couple stochastic simulations of genome evolution with a new supervised deep learning model to infer key parameters of molecular evolution. Our model is designed to directly analyze multiple sequence alignments and estimate per-site evolutionary rates and divergence, without requiring a known phylogenetic tree. The accuracy of our predictions matches that of likelihood-based phylogenetic inference, when rate heterogeneity follows a simple gamma distribution, but it strongly exceeds it under more complex patterns of rate variation, such as codon models. Our approach is highly scalable and can be efficiently applied to genomic data, as we show on a dataset of 26 million nucleotides from the clownfish clade. Our simulations also show that the per-site rates obtained by deep learning increase the likelihood of the true tree and could therefore lead to more accurate phylogenetic inference. We propose that future advancements in phylogenetic analysis will benefit from a semi-supervised learning approach that combines deep-learning estimation of substitution rates, which allows for more flexible models of rate variation, and probabilistic inference of the phylogenetic tree, which guarantees interpretability and a rigorous assessments of statistical support.

## Introduction

Since the seminal work by J. Felsenstein (1973) to infer phylogenetic trees by maximum likelihood, evolutionary models based on probabilistic approaches have been the central modeling framework in phylogenetics. This has led to a tremendous increase in our ability to infer evolutionary relationships, to investigate the dynamics of molecular evolution, to model the evolution of complex traits across lineages and to test evolutionary hypotheses to advance our understanding of the factors shaping the tree of life (Felsenstein, 2003; Lemey et al., 2009). While other methods based on genetic distances or parsimony criteria are still employed, for instance, to initialize phylogenetic tree inference or to provide a fast preliminary description of evolutionary processes, probabilistic approaches are widely seen as the best practice in the field.

One of the main challenges with the development of probabilistic models of evolution is to ensure that the parameters incorporated in the models are valid and identifiable when applied to real biological data. Yet, it is difficult, and in most cases even impossible, to experimentally generate data that would allow us to observe evolution in action and validate the estimates of the model parameters inferred from the outcome of such experiments. Indeed, while experimental evolution is applicable to some organisms with short generation times (e.g. for bacteria; Lenski, 2017), simultaneously capturing the evolutionary dynamics that result in genome evolution and the traits involved in adaptation, remains impossible for the most part. This means that validating evolutionary models is challenging and, when analyzing these evolutionary dynamics, we are inherently unable to compare our model estimates with a ground truth.

To overcome this limitation, most methods to infer evolutionary processes use simulations to assess the identifiability of model parameters and assess the robustness of the estimation. These simulations are synthetic realizations of evolutionary processes that are obtained through stochastic simulations. In a phylogenetic framework, we can use birth-death processes to generate phylogenetic trees and apply Markov models of nucleotide substitutions to simulate the evolution of a DNA sequence along the tree. Then, the same stochastic processes are typically used to also derive a likelihood-based model to estimate the generating parameter values from the simulation outcomes. For instance, we can derive the likelihood of a phylogenetic tree under a birth-death process and use it to estimate the speciation and extinction rates (Nee et al., 1994; Gernhard, 2008). We can also derive the likelihood of a DNA sequence alignment under a Markov process of evolution and use it to infer the underlying tree (Felsenstein, 1981). Comparisons between simulated and estimated parameter values (e.g., the true vs inferred phylogenetic tree) are then used to assess the accuracy of likelihood-based inference. This approach is routinely used for molecular evolution (e.g. Zaheri et al., 2014), phenotypic evolution of quantitative and discrete traits (e.g. Harmon et al., 2010; Maddison and FitzJohn, 2015), phylogenetic inference (e.g. Salamin et al., 2005), species diversification (Rabosky, 2006; Stadler, 2011), fossil preservation (Heath et al., 2014; Silvestro et al., 2019), biogeographic inference (Landis et al., 2013; Hauffe et al., 2022).

Despite its apparent circularity, this is a robust approach to validate the ability of a likelihood-based model to recover the parameters of the generative process correctly and can be used to verify their identifiability (e.g. Ree and Sanmartín, 2018; Silvestro et al., 2018; Louca and Pennell, 2020). Further, simulations generated while violating model assumptions can help quantifying the limits of our models and the conditions where the models will fail. A potential limitation of this use of simulations is that they tend to be oversimplified realizations of the biological process, which can impact our assessments of model accuracy (Nute et al., 2019).

Beside the use of likelihood-based models to infer the biological processes of interest, there is a growing interest in machine learning approaches to detect patterns associated with evolutionary processes. The use of Deep Learning (DL) has quickly expanded into a wealth of applications across scientific fields (e.g., Jumper et al., 2021). In evolutionary biology, deep neural networks have been proposed to infer speciation and extinction rates (Silvestro et al., 2020; Lambert et al., 2023), to study coevolution (see Sapoval et al., 2022, for a review), but also, for example, to infer phylogenetic trees using quartets (Zou et al., 2019; Suvorov et al., 2019; Kulikov et al., 2023), perform substitution model testing (Abadi et al., 2020) or place new samples on an existing phylogenetic tree (Jiang et al., 2022). However, the development of DL in phylogenetics and evolutionary models is restricted because training datasets are scarce or cannot be generated unless we restrict our focus to short-term evolutionary processes. Further, DL approaches often consist of (or are viewed as) over-parameterized black-box models that do not allow a direct interpretation of the parameters, contrary to probabilistic approaches (Sapoval et al., 2022).

Although probabilistic inference and DL can be seen as very different methodologies to analyse data, there are analogies in how model validation is performed. Indeed, generative models of evolution based on stochastic simulations can be coupled with supervised DL models, just like the same simulations are used to benchmark likelihood-based models. We show here that this use of DL models can complement likelihood-based approaches to estimate parameters that are interpretable and relevant for evolutionary models (Fig. 1).

**Figure 1:**
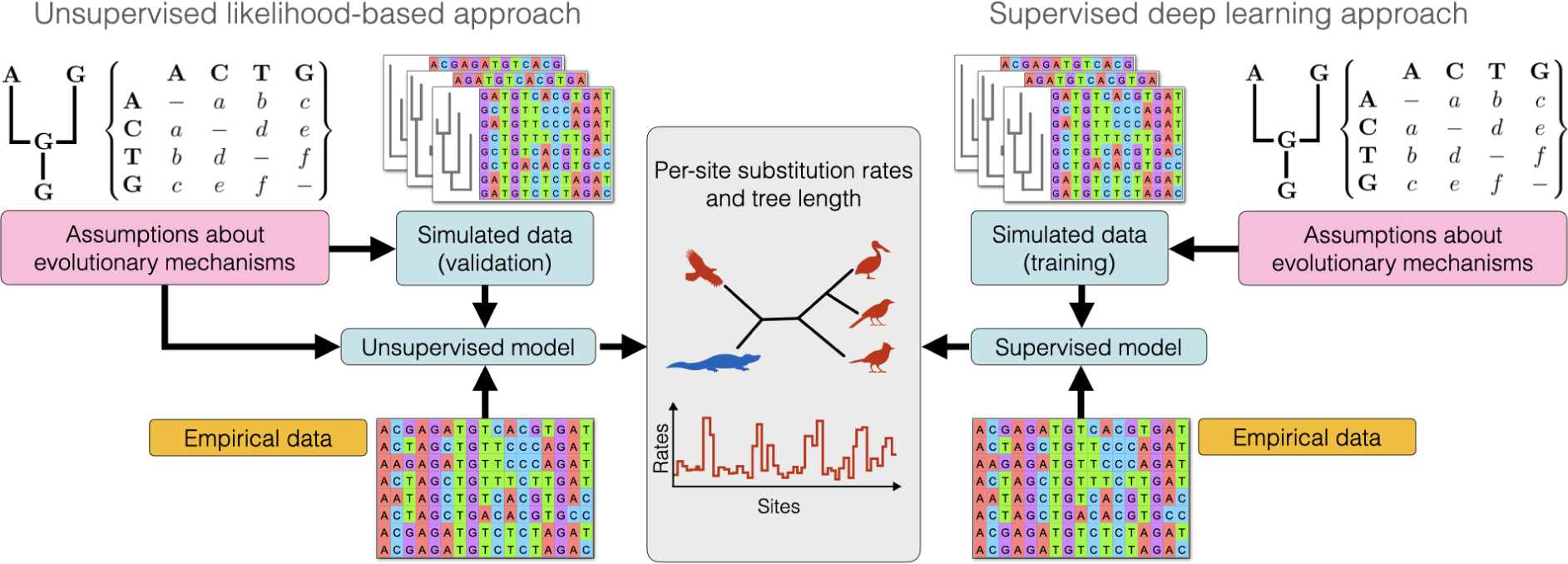
Schematic representation of the workflow used to estimate parameters of interest within an unsupervised model used within a likelihood framework (left) and a supervised model used with a DL framework (right). Both models use alignments of orthologous sequences of nucleotides (empirical or simulated data) to infer substitution rates and genetic distances. The simulations are used differently in the two cases, to either validate the model in a likelihood framework or to simulate training datasets in the DL framework.

In this paper, we develop a DL model to infer the rates of molecular evolution from a multiple species alignment of DNA sequences. We couple stochastic simulations with a new supervised learning model based on recurrent neural networks and sparse networks with parameter sharing. Our model and results show that the conceptual differences between standard unsupervised likelihood-based models and supervised DL are smaller than generally assumed in the context of molecular evolution. Predictions of site-specific substitution rates are robust across a range of evolutionary scenarios, with accuracy matching or exceeding that of state-of-the-art likelihood estimations. Our approach can efficiently analyze millions of sites to compare evolutionary rates at genomic scales. Finally, we show that the predicted rates have the potential to improve the likelihood-based estimation of phylogenetic trees, indicating that the application of this approach might have broader implications in phylogenetic inference.

Going forward, we propose that DL methods can be integrated within unsupervised likelihood approaches to help incorporating more realistic evolutionary scenarios in phylogenetic inference.

## Methods

### A deep learning model to infer molecular evolution

We developed a DL framework to estimate the total amount of divergence across a set of nucleotide sequences and site-specific substitution rates. The sequences are assumed to be part of an alignment of orthologous genes or genomic regions sampled across multiple species.

Our implementation includes two modules: a simulator that can efficiently generate realistic datasets of aligned nucleotide sequences and a DL module that can be trained on these datasets to make predictions from empirical data. Although we employed standard substitution models for generating nucleotide alignments, we incorporated a diverse array of modes of rate heterogeneity across sites, only a few of which are currently available in likelihood-based phylogenetic software. We used this to show that our framework can help identifying patterns that would otherwise be difficult to parameterize in a likelihood context.

### Simulating molecular evolution

We simulated the evolution of orthologous sequences within a phylogenetic framework assuming an independent Markov process of substitution at each site. We first generated a phylogenetic tree with a random topology and assigned exponentially distributed branch lengths sampled from *ν ∼* Exp(*λ*) with log(*λ*) *∼ u* (log(0.0002), log(0.2)). The total length of the phylogenetic tree was therefore *T* = ∑*_i_*(*ν_i_*), where *i ∈ {*1*,…*2*N −* 1*}* was the index of each branch in a tree of *N* species. We simulated the evolution of nucleotides based on three substitution models (JC, HKY, GTR; Jukes and Cantor, 1969; Hasegawa et al., 1985; Tavaré, 1986) using the program Seq-Gen (Rambaut and Grass, 1997) through its Python interface implemented in Dendropy (v.4.5.2 Sukumaran and Holder, 2010). Across simulations, we varied the model parameters, i.e. base frequencies and instantaneous substitution rates, by sampling them from distributions chosen to reflect a broad range of evolutionary scenarios (Table S1).

Since we focused our DL model on the inference of site-specific evolutionary rates, we implemented different distributions of rate heterogeneities across sites. Regardless of the mode of rate heterogeneity (Figs. S1, S2), site-specific evolutionary rates were always rescaled to relative rates, such that their mean across all sites equals 1 (Yang, 1994). First, we implemented a gamma mode, where site-specific relative rates were drawn from a gamma distribution, *r_i_∼* Γ(*α, β*) with the shape and rate parameters set equal and drawn from log(*α*) = log(*β*) *∼ u* (log(0.1), log(2)). This setting generated rates with an average value of 1, with increasing heterogeneity when the shape and rate parameters were small (Yang, 1993). This mode of rate heterogeneity reflects the standard gamma model of rate heterogeneity, which is however typically discretized in four or more rate classes (Yang, 1994), and which is almost ubiquitously used in phylogenetic inference. Second, we implemented a bimodal mode of rate heterogeneity where sites are randomly assigned a high or a low rate based on log(*r_i_*) *∼ {−m, m}* with *m* sampled from an exponential distribution Exp(1). Third, we implemented a spike-and-slab mode as a variation of the bimodal model, in which most sites evolve under low rates and few sites evolve under high rates. The low background rates were drawn from a log-normal distribution such that log(*r_i_*) *∼ N* (0, 0.1), while high rates were obtained by multiplying the background rates by a factor *m ∼ u* (2, 10). Sites were assigned to a high rate randomly with probability *r*, with log(*r*) *∼ u* (log(0.01), log(0.1)). Fourth, we simulated rates based on a non-stationary distribution obtained through a geometric Brownian motion process. In this case, a vector of rates was sampled from a geometric Brownian process such that log(*r_i_*_+1_) *∼ N* (log(*r_i_*)*, σ*), with *σ ∼ u* (0.02, 0.2). Fifth, we implemented a codon mode of rate heterogeneity, in which triplets of nucleotides were assigned low, very low, and high rates, for the first, second and third positions, respectively (Nielsen and Yang, 1998). We sampled the rate of the second position from a log-normal distribution, such that for triplet *i*, 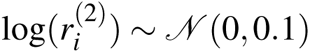. Rates for the first and third positions were then obtained as 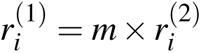, with *m ∼ u* (1, 5) and 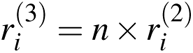, with *n ∼ u* (5, 15), respectively.

We additionally simulated datasets with rates varying among a variable number of blocks of adjacent sites, thus introducing auto-correlation in rate heterogeneity (Fig. S2). We drew the number of blocks from a geometric distribution with a mean of 100 and truncated at 1000 (i.e. the number of sites in the alignments), and randomly sampled the sizes of the blocks. We then applied rate variation among blocks using the gamma, bimodal, spike-and-slab and geometric Brownian modes described above. Finally, we included datasets generated under mixed modes of rate heterogeneity (Fig. S2), in which the alignment was split into two blocks of random size, each with its own randomly selected rate heterogeneity mode.

### Architecture and training of the deep learning model

We implemented a DL model (hereafter called phyloRNN) that takes an alignment of nucleotides as input and returns two outputs: site-specific relative rates of evolution and the expected number of substitutions per site, which is equivalent to the sum of all branch lengths in a phylogenetic inference framework. For simplicity, hereafter we will refer to this second output as *total tree length*, even though our model does not use any tree representation. We built our model based on a recurrent neural network (RNN) to capture the sequential nature of the input data. Specifically, we used a bidirectional long short-term memory architecture (bLSTM; Hochreiter and Schmidhuber, 1997; Graves and Schmidhuber, 2005; Gers et al., 2000), with a site-specific output, which is a multidimensional representation of the initial alignment. Each site-specific output of the bLSTM layer fed into two individual fully connected deep neural networks with parameters shared across all sites. The first network returned a site-specific relative rate. The output of the second network was instead concatenated across all sites and fed into a fully connected layer to return a prediction of the total tree length. A schematic representation of the model is shown in Fig. S3.

After preliminary testing based on validation accuracy, we chose the following architecture for our experiments: two bLSTM layers of 128 and 64 nodes, respectively, with a tanh activation function and sigmoid recurrent activation. The output of the second bLSTM layer, for each alignment, was thus of shape (n. sites *×* 64) and served as input to the two deep networks that output site-specific rates and tree length. For the site-specific rates, we used a neural network with two hidden layers of 64 and 32 nodes and a swish activation function (Ramachandran et al., 2017), followed by an output layer with 1 node and a softplus output function (Szandała, 2021), reflecting a distribution of rates constrained to positive values. The concatenated output (one value for each site), was then subjected to a rescaling function dividing each value by the mean across all sites. This rescaling ensures that the estimated rates had a mean equal to 1, thus transforming them into relative rates. The output of the second bLSTM layer was also used to infer the log-transformed total tree length, by feeding into site-specific deep networks with two hidden layers of 64 and 1 nodes and swish activation functions. As for the site-specific rates all networks shared the same parameter values. Their output (of size 1 for each site) was then concatenated across sites and fed into a fully connected hidden layer with 8 nodes and swish activation function. The output layer had one node and a linear activation function reflecting the negative-to-positive plausible range of values for the log-transformed tree length.

We trained our model based on 10,000 simulated alignments of 50 species and 1,000 sites, each with a randomly selected mode of rate heterogeneity, of which 80% were used as a training and 20% as validation set. We used the same phyloRNN model trained on an equal mix of simulated datasets for all the comparisons outlined below. Each alignment was transformed into a series of two-dimensional arrays of numbers after one-hot-encoding the sequence of nucleotides. The data fed to the model was thus composed of 1,000 two-dimensional arrays (one per site in the alignment) with each a size defined by the number of species times the one-hot encoding of each nucleotide present at a single site (i.e. 50 species *×* 4 states = 200). These two-dimensional arrays were then stacked on a third dimension representing the number of instances (i.e. batches used in the training or datasets used for testing) analysed to create the final input for our model (n. instances, n. sites, n. species *×* n. bases).

The training and validation losses were calculated as the sum of the mean squared errors (MSE) computed across per-site rates and total tree length. We log-transformed tree lengths to reduce the range of loss values and improve the efficiency of the optimization. We trained the model over multiple epochs with a batch size of 100 and monitoring the validation loss with a patience parameter set to 20 epochs. The model was calibrated in fewer than 100 epochs, after reaching the lowest validation accuracy combining the loss functions assigned to the per-site relative substitution rates and the log-transformed tree length, (Fig. S4). We kept the model parameters inferred from the epoch with lowest validation loss. We implemented our model based on the Functional API of the Tensorflow module (Abadi et al., 2015) and trained it using the RMSprop optimizer with learning rate set to 1*e−* 3. Our phyloRNN model and the associated scripts are available in a open-source repository: github.com/phyloRNN.

### Accuracy of rate and tree length estimates

We validated the performance of our model based on a test set of 600 alignments simulated under the different modes of rate heterogeneity described above. We compared the accuracy of the estimated per-site relative rates and number of substitutions based on our phyloRNN model with those obtained through maximum likelihood phylogenetic inference. Specifically, we analyzed each alignment with PhyML v.3.3 (Guindon et al., 2010) running a maximum likelihood optimization. In a typical phylogenetic analysis, model testing is carried out first to select the best fitting substitution model (Abadi et al., 2019). Here, we assumed that the true substitution model (here one among JC, HKY, and GTR; Table S1) was known and we used it in the PhyML optimization to reduce computing time. We repeated the phylogenetic analyses under two rate heterogeneity models: i) the discrete gamma model (Yang, 1994) –by far the most commonly used in phylogenetic inference– and ii) the more flexible –but less frequently used– free-rates model, which allows for a number of rates to be inferred without making a specific assumption about their distribution (Soubrier et al., 2012). We obtained the marginal per-site rates using the –print_site_lnl command in PhyML (column Posterior mean in the output file). We quantified the average number of substitutions per site as the sum of all branch lengths from the inferred phylogenetic tree.

After obtaining the relative rates and average number of substitutions per site under both phyloRNN and the two likelihood models (discrete gamma and free-rates), we compared them against the respective true values to quantify their accuracy. We used MSE across all sites and all simulations to quantify the performance of the different models and additionally computed the coefficient of determination R^2^ for each alignment to compare the estimated pattern of rate variation across sites against the true rates. To further explore the results, we divided the test set into subsets based on the rate heterogeneity model that was used to generate them (Figures S1, S2) and calculated MSE and R^2^ for each subset. We calculated the same summary statistics to evaluate the accuracy of the estimated tree length.

### Impact of rate estimated by phyloRNN on tree inference

We evaluated the potential effects of using `phyloRNN` estimates of rate heterogeneity in phylogenetic inference. Since standard phylogenetic software does not allow tree inference with predefined per-site rates, we used two indirect approaches to approximate the impact of using `phyloRNN` rates as opposed to the two existing models implemented in `PhyML` (discrete gamma and free-rates).

First, we compared the likelihood obtained on the true tree using the estimated rates and the true simulated per-site rates. We computed the likelihood while fixing the topology and branch lengths to their true values (i.e., the simulated tree) and using 1) the true simulated per-site rates, 2) the marginal per-site rates estimated by PhyML under the discrete gamma and free-rates models, and 3) the per-site rates estimated with phyloRNN. We recomputed the likelihood of the true tree based on the fixed rates per site and the true parameters for the substitution model (script available at github.com/phyloRNN/phyloRNN). For each simulated alignment, we compared the likelihood obtained by the true versus estimated rates by calculating the difference in log-likelihood. We summarized these comparisons by computing the proportion of datasets in which the likelihood of the true tree substantially decreased (i.e. more than 2 log-likelihood unit differences) using estimated versus true rates and interpreted them as a measure of the potential impact of the gamma, free-rates, and phyloRNN models on the accuracy of tree inference.

Second, we compared, for each model used to estimate rates per site, the likelihood of the true tree against the likelihoods of a posterior sample of trees obtained from a Bayesian analysis. To this aim, we first obtained a posterior sample of 50 trees for each test dataset using MrBayes (Ronquist et al., 2012) under a GTR + gamma model and default prior settings. We then recomputed the likelihood of both the set of sampled trees and the true tree based on the rates inferred under the gamma and free-rates models (as inferred with PhyML) and under the phyloRNN model. Finally, we ranked all trees by their likelihood to assess whether the likelihood of the true tree falls within the ranged of sampled values in the posterior set of trees.

If the estimated rates adequately reflected the true rate variation in the data, we expected the likelihood of the true tree to rank somewhere within the range of likelihoods of the sampled trees, as this would indicate that the true tree is likely to be included in the posterior distribution obtained under a gamma model. In contrast, if in a substantial proportion of simulations the likelihood of the true tree was lower than that of the sampled trees, then the estimated rates might be inadequate by assigning significantly higher likelihood to solutions that differ from the truth. Finally, if in a substantial proportion of simulations the likelihood of the true tree is higher than that of the sampled trees, then the rates, e.g. estimated through a free-rates or phyloRNN model, would favor sampling the true tree over alternatives sampled under a discrete gamma model.

Since the trees were obtained under a gamma model, we expected the likelihood of the true tree based on rates from the gamma model to be found within the sampled range or lower (if the rates are inadequate). A higher likelihood is still possible, however, if the subset of 50 trees does not capture the actual upper boundary of the posterior distribution of likelihood values. With rates inferred from the free-rates or phyloRNN models, we instead expected a larger proportion of simulations in which the true tree returned a higher likelihood than the sampled ones because of the better fit to the underlying true mode of rate heterogeneity.

### Analysis of the clownfish genomes

We applied our model to a genomic dataset of 28 species of clownfish representing the first chromosome (Marcionetti and Salamin, 2023). Even though phyloRNN does allow for gaps in an alignment, we decided for simplicity to filter out positions in the alignment containing gaps and ambiguities reducing the initial dataset of over 46 millions nucleotides to a total of 26,294,222 aligned nucleotides. We trained a new model containing 10,000 simulated datasets to match the input of 28 taxa and 1,000 nucleotides. We used it to predict per-site relative substitution rates and total tree length (as a measure of overall divergence across the chromosome) through non-overlapping sliding windows of 1,000 sites. We generated histograms of the rate heterogeneity across a random subset of exons (filtering out those of length smaller than 500 nucleotides) to visually assess whether a gamma distribution adequately approximates the empirical distribution of rates. Finally, we tested whether protein-coding regions showed consistently lower substitution rates compared with neighboring non-coding regions as expected if they are functionally constrained. We estimated the mean substitution rate for each exon (filtering out those of length smaller than 250 nucleotides; results did not change if the limit was set to 100 nucleotides) as well as the mean rates of the directly adjacent non-coding regions selecting 250 nucleotides before and after the start or end of each exon selected. We performed paired t-tests to test whether the rates were significantly different between exons and directly adjacent regions.

## Results

### **Performance of the** phyloRNN **model**

We measured MSE and R^2^ values for the rate predictions obtained from the trained model on the test set and the rates estimated by PhyML with the gamma and free-rates models. The results showed that our model provided substantially more accurate estimations of the per-site rates compared with maximum likelihood estimates obtained through a gamma model of rate heterogeneity (lower MSE and higher R^2^ values; Table 1). The phyloRNN model also outperformed maximum likelihood estimations based on the more flexible free-rates model under most scenarios, especially the more complex heterogeneity modes (Table 1), although the difference was smaller than with the gamma model. After breaking down the results by the simulated mode of rate heterogeneity, we found that the improvement in rate estimation is particularly strong in the case of codon mode of rate heterogeneity (MSE values decreasing by one order of magnitude when using phyloRNN) and in the case of autocorrelated rates (Fig. 2). The phyloRNN estimates appeared to consistently outperform maximum likelihood estimates particularly in their ability to recover multimodal distributions of rates across sites (Figs. 2, S5–S7).

**Figure 2:**
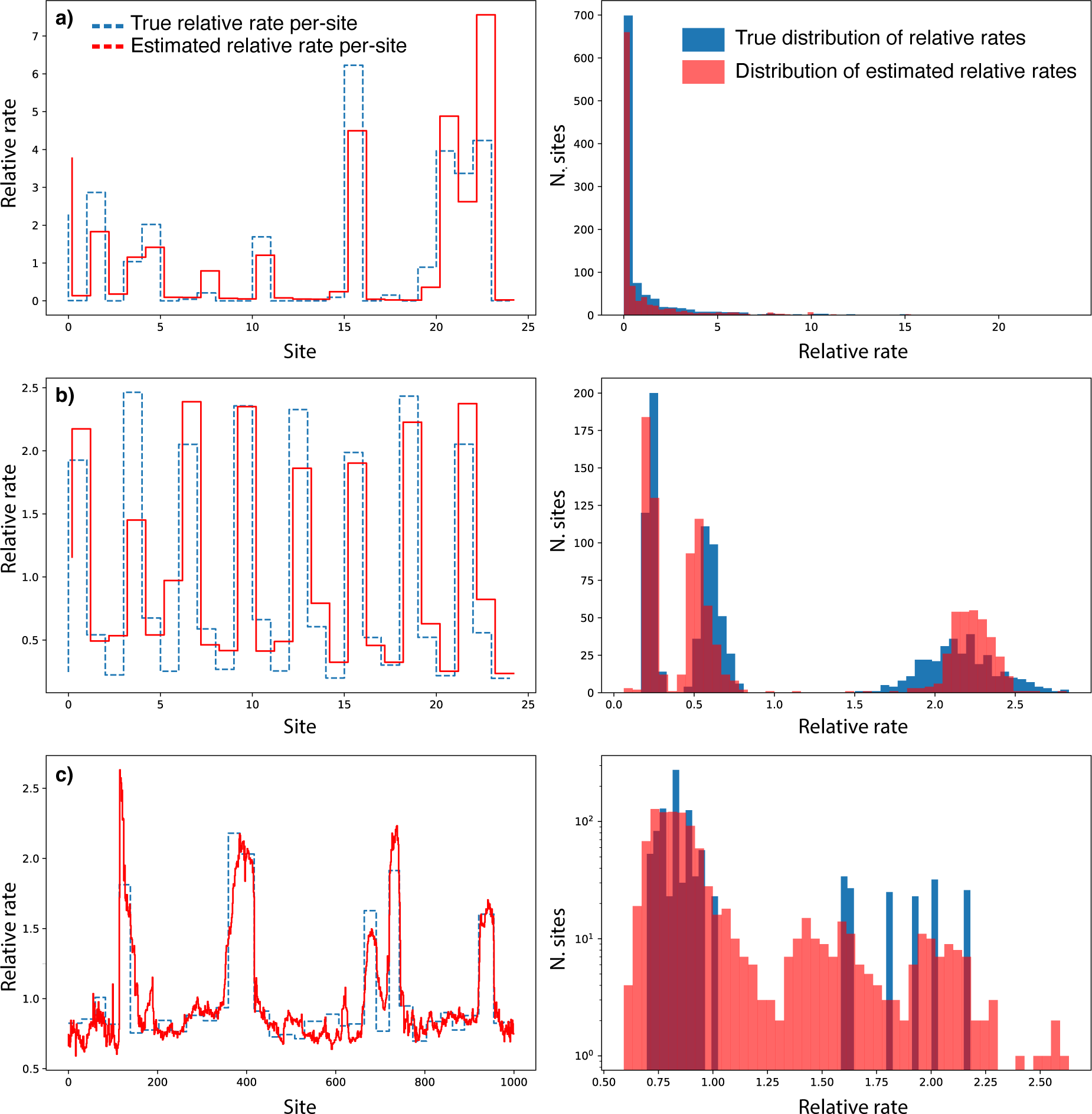
Example of simulated and estimated per-site rates based on our DL model. Plots on the left show per-site rates (note that the estimated rates are shifted slightly to the right for clarity). Histograms show the true distribution of rates across 1000 sites in an alignment (the bottom right is in log space for clarity) and the distribution of estimated rates. The simulations show an example of different modes of rate heterogeneity: gamma (a), codon (b), and spike-and-slab autocorrelated (c).

**Table 1:** Accuracy of site-specific rates estimated under different models calculated across a test set of 600 datasets generated under different modes of rate heterogeneity. The accuracy is quantified a mean squared error and as R^2^ (in parenthesis), comparing the true rates with those predicted through two maximum likelihood models (gamma and free-rates) and through DL (phyloRNN). DL rate estimates consistently outperform those from a gamma model, in most cases matching or slightly exceeding the accuracy of the free-rates model. DL substantially outperforms likelihood methods in simulations with autocorrelated rates or based on codon models.

| Mode of rate heterogeneity | Gamma model | Free-rates | phyloRNN |
| --- | --- | --- | --- |
| All combined | 0.833 (0.175) | 0.705 (0.206) | 0.614 (0.347) |
| Gamma | 1.869 (0.409) | 1.311 (0.598) | 1.64 (0.446) |
| Bimodal | 0.454 (0.143) | 0.28 (0.139) | 0.332 (0.063) |
| Spike-and-slab | 0.538 (0.161) | 0.32 (0.484) | 0.517 (0.08) |
| Geometric Brownian | 1.469 (0.337) | 1.408 (0.387) | 1.387 (0.268) |
| Codon | 0.379 (0.341) | 0.405 (0.316) | 0.076 (0.942) |
| Gamma autocorrelated | 1.578 (0.465) | 1.613 (0.584) | 0.755 (0.798) |
| Bimodal autocorrelated | 0.329 (0.133) | 0.25 (0.112) | 0.178 (0.294) |
| Spike-and-slab autocorrelated | 0.387 (0.032) | 0.323 (0.03) | 0.271 (0.109) |
| Geometric Brownian autocorrelated | 0.276 (0.034) | 0.27 (0.028) | 0.113 (0.457) |
| Mixed | 1.882 (0) | 1.656 (0) | 1.689 (0) |

Although the phyloRNN model did not use (or attempt to estimate) a phylogenetic tree, its estimation of the total tree length is unbiased (Fig. 3) and showed comparable accuracy with the estimates obtained from a maximum likelihood analysis based on a gamma model of rate heterogeneity, while the free-rates model generally produced the most accurate estimations. The accuracy of tree length estimation was high in most simulations with mean absolute percentage errors generally below 15% (Table 2). Tree lengths were inferred with higher error for simulations based on mixed modes of rate heterogeneity and this was the case across all three inference methods.

**Figure 3:**
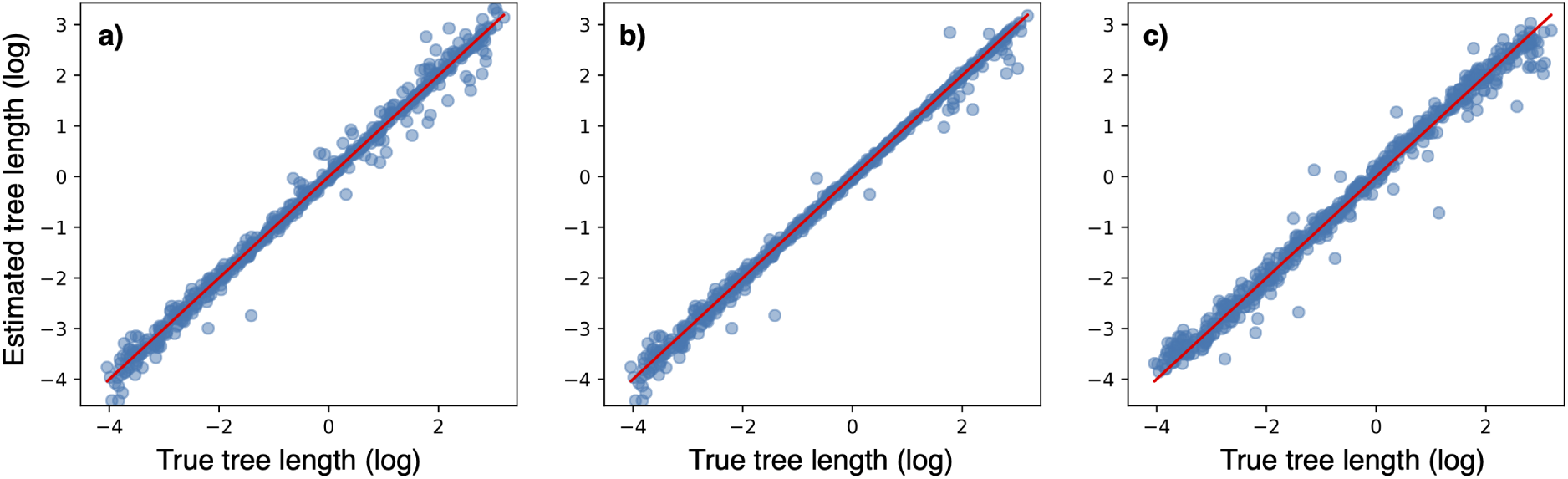
Estimated total tree length (log-transformed) under maximum likelihood models assuming a gamma heterogeneity model (a) and a free-rates model (b), and through our DL model (c). The results are shown for a test set of 600 datasets generated under different modes of rate heterogeneity (more details in Table 2).

**Table 2:** Accuracy of total tree length estimated under different models calculated across a test set of 600 datasets generated under different modes of rate heterogeneity. The accuracy is quantified a mean absolute percentage error and as R^2^ (in parentheses), comparing the true rates with those predicted through two maximum likelihood models (gamma and free-rates) and through DL (phyloRNN). The accuracy of DL tree length estimates is generally similar to that obtained from maximum likelihood methods, notably outperforming the gamma model when the underlying data are simulated with gamma-distributed rate heterogeneity model. Maximum likelihood inference outperforms DL in simulations based on bimodal, codon, and geometric Brownian autocorrelated models.

| Heterogeneity model | Gamma model | Free-rates | phyloRNN |
| --- | --- | --- | --- |
| All combined | 0.106 (0.933) | 0.075 (0.955) | 0.152 (0.902) |
| Gamma | 0.147 (0.977) | 0.083 (0.987) | 0.158 (0.902) |
| Bimodal | 0.105 (0.936) | 0.047 (0.999) | 0.115 (0.953) |
| Spike-and-slab | 0.066 (0.993) | 0.055 (0.998) | 0.117 (0.986) |
| Geometric Brownian | 0.118 (0.955) | 0.102 (0.929) | 0.150 (0.934) |
| Codon | 0.056 (0.999) | 0.051 (0.999) | 0.103 (0.971) |
| Gamma autocorrelated | 0.156 (0.952) | 0.085 (0.921) | 0.173 (0.791) |
| Bimodal autocorrelated | 0.091 (0.925) | 0.057 (0.996) | 0.161 (0.957) |
| Spike-and-slab autocorrelated | 0.057 (0.995) | 0.053 (0.976) | 0.103 (0.971) |
| Geometric Brownian autocorrelated | 0.055 (0.999) | 0.052 (0.999) | 0.177 (0.98) |
| Mixed | 0.471 (0.605) | 0.459 (0.681) | 0.493 (0.60) |

### Impact on tree estimation

In our simulations, the likelihood of the true tree based on the true per-site rates substantially exceeded (Δlog *L >* 2) the likelihood of the same tree using a discrete gamma model of rate heterogeneity in 90.2% of the datasets. This means that, in a large fraction of the simulations, a gamma model decreased the likelihood of the correct underlying phylogenetic tree (Table S2). We observed a similar outcome under the free-rates model, where the likelihood of the true tree decreased in 89.6% of the simulations. In contrast, phyloRNN rates resulted in a substantially lower likelihood for the true tree only in 29.7% of the simulations, suggesting that the use of these rates in phylogenetic inference might result in more accurate estimated trees (Table S2).

A change in absolute likelihood does not necessarily imply that a model is less likely to sample the true tree in phylogenetic inference, because it could simply reflect a homogeneous shift in the likelihood surface. We therefore also evaluated the ranking of the true tree within a posterior sample of trees for each testing dataset described above. We sampled 50 trees from the posterior samples of MrBayes and ranked them in decreasing order based on their likelihood recomputed under predicted rates from the gamma, free-rates and phyloRNN models as above. We then compared the likelihood of the true tree obtained with the same models with the ranked sampled trees. Under the gamma model, the likelihood of the true tree was found within the range of sampled trees in 86.0% of the simulations and ranked first in 1.8% of the cases. In the latter case, the mean difference in log-likelihood between the true tree and the best tree from the posterior samples was small (ranging from −0.130 to −7.757, median of −1.857), which suggests that the true tree would in fact be included in a more extensive posterior sample including more than the 50 trees considered here. However, the true tree ranked last, and thus was outside the sampled range, in 12.2% of the simulations, with a range of log-likelihood differences between the true tree and the worst tree from the posterior samples ranging from 0.146 to 2129.506 (median of 24.137). This indicates that, for most of these cases, the true tree was unlikely to be sampled by the Bayesian algorithm, and thus excluded from the estimated posterior distribution of trees.

In contrast, under the rates estimated with phyloRNN, the true tree ranked last in only 3.7% of the simulations (log-likelihood difference ranging from 0.309 to 3602.207 with a median of 16.141), while it ranked first in 13.5% (log-likelihood difference ranging from −0.006 to −1841.836 with a median of −38.206). This suggests that in a substantial proportion of datasets, the phyloRNN rates would favor the true tree over the trees sampled under a gamma model. For comparison, the free-rates model performed similarly to phyloRNN, with 3.2% of simulations with the true tree ranking last (log-likelihood difference ranging from 0.033 to 2806.083 with a median of 10.348) and 12.2% of the simulations with the true tree ranking first (log-likelihood difference ranging from −0.125 to −1341.208 with a median of −20.146).

The rank of the true tree in all these comparisons did not depend strongly on the mode of rate heterogeneity, the model of substitution or the tree length used in the simulations which were not significant when analysed with a linear model. However, we did find that instance where the true tree ranked last under a gamma model (therefore being excluded from the sampled posterior distribution of tree) were associated with significantly higher error in both estimated per-site rates and tree length (Fig. S8). This corroborates the idea that an improved estimation of these parameters can lead to a more accurate tree estimation.

### Evolutionary rates and divergence in clownfish

The estimated substitution rates along chromosome 1 in clownfish showed a substantial degree of heterogeneity across sites, with 99% of the values ranging between 0.09 and 0.69, thus encompassing a 7-fold rate variation (Fig. 4 a). The overall distribution of rate heterogeneity across all *∼*26M sites followed quite closely a gamma distribution (Fig. 4 a), indicating that a gamma model should approximate well the true rate variation at a broad genomic scale. However, the distributions of substitution rates across smaller genomic regions showed that across-site rate heterogeneity often diverged substantially from a gamma distribution, showing multimodal patterns and heavy-tailed distributions (Fig. 4 b-i).

**Figure 4:**
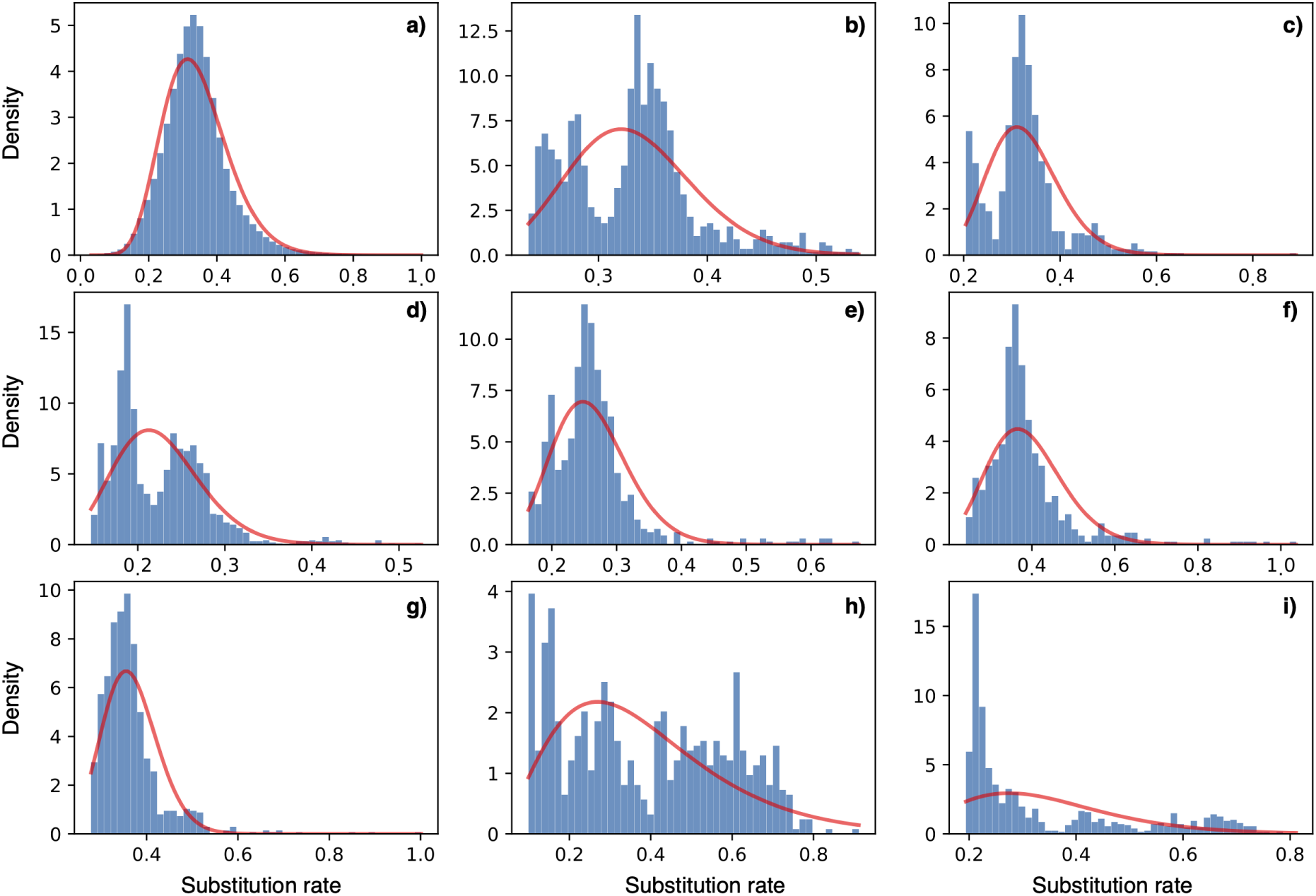
Estimated rate heterogeneity across sites plotted for all sites in chromosome 1 in the clownfish genome (a) and across a random sample of eight exons (b–i). The red lines show gamma distributions fitted to the data shown in the histogram. The rates match well a gamma distribution of rate heterogeneity across all 26+ million sites (a), consistently to the heterogeneity models typically used in likelihood-based phylogenetic analyses. However, at the exon level, the rate distribution often diverges substantially from that of a gamma distribution, displaying multimodal or heavy-tailed distributions. For improved visualization, rates > 1 (N = 34,085, or *∼*0.12%) are not shown in panel a).

The degree of clownfish divergence (the total tree length) estimated within blocks of 1,000 sites revealed up to *∼*4-fold variation in estimated number of substitutions across the chromosome. The distribution of substitution rates per site also highlighted regions in the chromosome spanning several thousands of sites that are more conserved (i.e. low average rates) and others characterized by much higher rates (Fig. 5).

**Figure 5:**
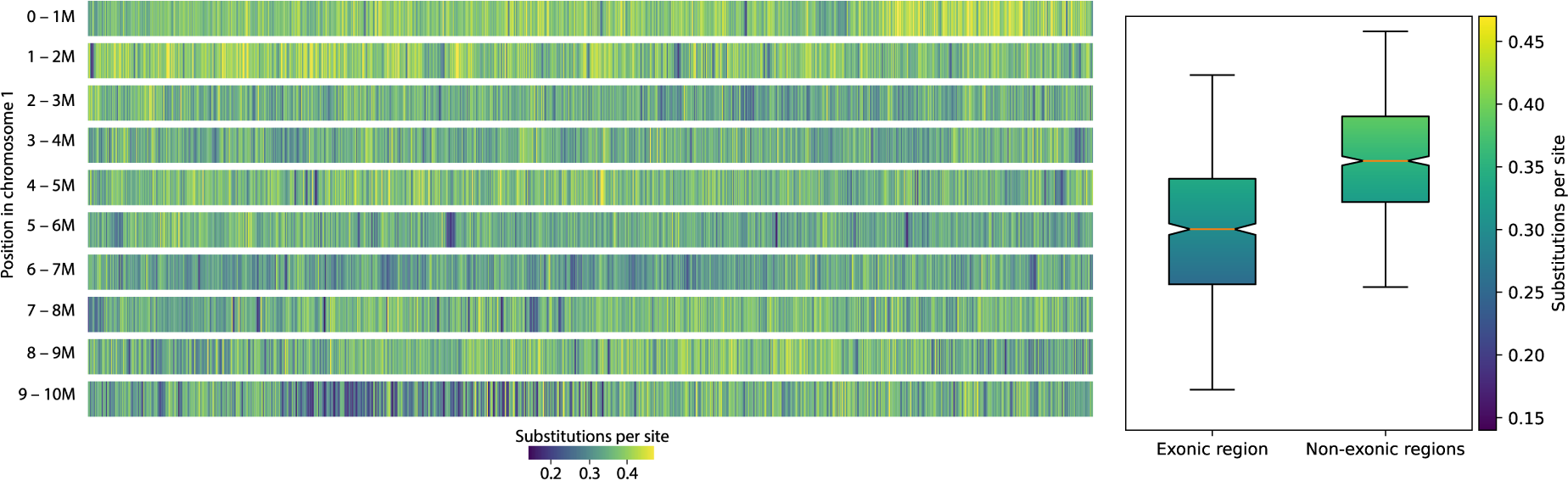
Estimated number of substitutions per site inferred through our RNN model across the first 10 million nucleotides of the clownfish genomes (28 species, chromosome 1). Left panel: Substitutions per site as function of chromosomic position, rates are averaged across blocks of 1,000 nucleotides. Right panel: Box-plots of substitutions per site for exonic region (random sample of 854 exons, minimum of 250 nucleotides) and adjacent non-exonic regions (250 nucleotides before and after the start or end of each exon).

When comparing the substitution rates between coding and non-coding regions, we found substitution rates in exons to be lower than the rates in adjacent regions in the chromosome in 82% of the 854 exons analyzed. The mean rate in exons was on average *∼*14% lower than in the adjacent non-coding regions (Fig. 5). Paired t-tests showed that this rate difference was overall significant (p-value *<* 1*e^−^*^90^*, T < −*23.00, 95% confidence interval: [-0.06, −0.05]), while the left and right adjacent regions did not differ from one another (p-value = 0.26*, T < −*1.12, 95% confidence interval: [-0.01, 0]).

## Discussion

### A new deep learning model to infer molecular evolutionary rates

We presented a new framework using stochastic simulations to train a DL model and estimate site-specific rates of evolution and total tree length from an alignment of nucleotide sequences. We specifically designed the architecture of our model to reflect the characteristics of molecular data and assumptions behind the evolution of DNA sequences. Indeed, the bLSTM layers capture the sequential nature of DNA data, while the use of site-specific networks with parameter sharing reflects the fact that each feature in the input layer is an instance of the same nature (i.e. a nucleotide).

Our implementation allowed us to estimate with high accuracy the evolutionary rates for each site of an alignment together with the total evolutionary divergence in the alignment, which, in a phylogenetic context, is quantified by the tree length. This was done using solely the information from the alignment and without the addition of a defined phylogenetic tree in the input to our phyloRNN model. We showed that our model outperforms likelihood estimations based on the standard gamma distribution of rate heterogeneity and that it matched or outperformed, depending on the rate heterogeneity mode used, the estimates from the more parameter-rich and less frequently used free-rates model. The computational efficiency of our approach makes it highly scalable, allowing for the analysis of large-scale genomic data.

### Evolutionary inference using phyloRNN

While rate heterogeneity is generally considered as a nuissance parameter in phylogenetic inference (Yang, 1994), an accurate estimation of substitution rates per site can be used to identify genomic regions that are under evolutionary constraints or deviate from the average pattern across genomic regions (Mayrose et al., 2005). It can also be used to identify protein coding genes because of the significantly reduced rate of evolution compared with adjacent genomic regions, as we found in the clownfish dataset (Fig. 5). These results could be coupled with other approaches used in genomics to help annotate *de novo* genome assemblies or to identify regions of interest that show unexpected levels of conservation outside of the protein-coding genes.

The analyses of the genomic data for clownfish allowed us to look at the distribution of rates across a large empirical dataset. Although the distribution of rates corresponds to a gamma distribution when the full set of ca. 26 million sites are considered, the distributions for individual exons can drastically differ from a gamma distribution (Fig. 4). A similar pattern had been previously shown in the distribution of average rates across genes (Bevan et al., 2007) instead of individual sites like in our study, or in datasets with low level of variation (Jia et al., 2014). To tackle such cases in a likelihood framework, complex mixture of gamma distributions can be used to model this distribution of rate heterogeneity (Mayrose et al., 2005), but at the cost of additional complexity during the optimization process (Bevan et al., 2007). In contrast, our phyloRNN model can easily account for complex distributions through the simulation of various types of rate heterogeneity.

The gamma distribution to model rate heterogeneity was introduced in a landmark paper by Z. Yang in 1994 (Yang, 1994) and led to a substantial improvement in likelihood-based tree inference across most empirical datasets. Although the fit of models of evolution including a gamma distribution is drastically improved when compared to models without rate heterogeneity, the use of a gamma distribution is not based on a biological assumption. Additionally, our results showed that deviations from the gamma distribution can have a large impact on the basic estimates of rates per site or tree length (Tables 1, 2). Alternative models have been proposed to include more biological realism (Heaps et al., 2020), and allow for more flexible distributions, like the free-rates model which is implemented in some phylogenetic software (e.g., the PhyML program used here and IQTREE (Minh et al., 2020)). Other programs use a discrete-rates CAT model, implemented in RAxML (Stamatakis, 2014) and FastTree2 (Price et al., 2010), which is more efficient computationally than fitting a gamma distribution. Finally a full Bayesian method to simultaneously estimate the substitution model and rate at each site has also been proposed (Wu et al., 2013), but it involves a large number of parameters with unclear effects on tree inference. The model we developed here could lead to an alternative, potentially more efficient, approach, where rates per site are inferred prior to phylogenetic inference and without assuming any specific distribution. This approach would reduce the number of free parameters in the phylogenetic inference model and potentially have a beneficial impact on its accuracy.

### Impact on phylogenetic inference

Including rate heterogeneity has been shown to have a large impact on phylogenetic inference (Yang, 1994; Sullivan and Swofford, 1997; Abadi et al., 2019). Our simulations further demonstrated that an accurate modeling of the rate variation across site will also affect, sometimes drastically, the estimation of the tree likelihood. The effect is not simply a monotonic increase or decrease of the likelihood surface, but shows also an impact on the ranking of the trees. We showed it indirectly by comparing the likelihood of sampled trees computed by assigning rates per sites estimated using the gamma, free-rates and phyloRNN models. The changes in log-likelihood values demonstrate that an inaccurate estimation of the rates per site can bias tree inference and result in significantly lower likelihood of the true tree compared to alternative hypotheses.

In likelihood-based phylogenetic inference, nucleotide frequencies are routinely set equal to their empirical values calculated from the alignment, rather then estimated in the analysis. This is done to reduce the number of free parameters in likelihood optimization or posterior sampling algorithms. Similarly, per-site rates could be estimated from the alignment using our phyloRNN model before fixing them during the likelihood search. Flexible and modular phylogenetic programs like RevBayes (Höhna et al., 2016) or BEAST (Bouckaert et al., 2019) could easily accommodate rates obtained independently *a priori* within their tree search algorithms or even include a DL rate estimation as the first step to phylogenetic inference with limited additional computational burden.

### Performance of phyloRNN on big data

Our application of the phyloRNN model to chromosome 1 of 28 clownfish species showed that a trained model can rapidly estimate per-site substitution rates across large genomic datasets with a small computational footprint. The current implementation supports CPU parallelization of the simulations required for model training and allows for the generation of new models for the size of the datasets that we used in this study within few hours on a 64-CPU cluster. The use of GPU computing for model training is likely to further accelerate both training and prediction tasks (Abadi et al., 2015). On a standard laptop-grade CPU and without parallelization, we obtained predictions across 26 million sites in *≈*20 minutes, requiring less than 10 Gb of RAM. Performing these estimations with a standard phylogenetic software would be much more challenging both in terms of memory usage and CPU time. Our tests on a subset of the clownfish genomic data using PhyML indicate that a comparable analysis uses up to several tens of Gb of RAM depending on the number of site patterns present in the alignment and exceeds 40 hours, even with a fixed tree topology and optimization limited to branch lengths and model parameters.

### Effects of violations of model assumptions

A common critique to supervised DL models over their unsupervised likelihood-based alternatives in regression and other inference tasks, is their unpredictably erroneous behavior when presented with data that differ from the training data (Marcus, 2018). In the case in which the training data were simulated under a generative model, like in our study, differences between training and empirical data could be driven by violations of the assumptions of the generative model in real world evolution. However, violations of the model assumptions have also been shown to lead to wrong estimations in likelihood-based inference of evolutionary models. For instance, simplistic substitution models assuming equal substitution rates among nucleotides (an assumption clearly violated by the real evolutionary process), have long been known to lead to wrong tree topologies (D’Erchia et al., 1996; Sullivan and Swofford, 1997). More recently, Meyer et al. (2019) found that the presence of co-evolving sites in an alignment, which violates the common assumption of site independence, can bias in unpredictable ways phylogenetic inference, affecting the accuracy of both tree topology and branch lengths. Similar misbehavior in likelihood-based inference has been shown in the context of models of trait evolution Duchen et al. (2021) and species diversification (Louca and Pennell, 2020). Thus, phylogenetic and macroevolutionary analyses are likely to be generally sensitive to model violations.

Recent research has shown that the current models of nucleotide evolution might be inadequately reproducing realistic nucleotide sequence alignments (Trost et al., 2023), although the effects of this inadequacy on phylogenetic inference remains to be fully explored. While in likelihood-based models the assumptions about how evolutionary mechanisms play out are built directly into the likelihood function itself, in our phyloRNN framework the same assumptions are encoded in the simulation module (Fig. 1). This architectural difference makes it substantially easier to relax these assumptions in a model like phyloRNN that couples stochastic simulations with DL. We have demonstrated this through the implementation and training of a single model able to account for a range of heterogeneity patterns, including auto-correlated rates and codon models, each of which would require a specific parameterization in a likelihood framework. In the phyloRNN framework, the inclusion of additional heterogeneity patterns is straightforward as long as such patterns can be simulated, thus facilitating its extension to more diverse evolutionary scenarios.

### Why we still need likelihood-based evolutionary models

While DL is now permeating many research fields in biology (Sapoval et al., 2022), we think that well-principled and fully interpretable likelihood models will continue to play a key role in evolutionary biology, for several reasons. First, likelihood-based methods are (arguably) more suitable for the estimation of complex parameters. In contrast, most of DL models are designed to infer simple output parameters (e.g., continuous values in regression tasks or categorical variables in classification tasks). Their application to more complex parameters such as the phylogenetic tree topology is instead less straightforward to implement in a standard output layer beyond small scale implementations (e.g., Zou et al., 2019; Sapoval et al., 2022). Additionally, although we showed that a single DL model can be used to jointly infer different parameters, supervised models will typically not benefit from a joint parameter estimation (Marcus, 2018), unlike likelihood-based, and especially hierarchical Bayesian models (Gelman et al., 2013).

Second, likelihood-based methods provide a more direct and robust assessment of parameter uncertainty, e.g. through bootstrap values, confidence interval estimates or posterior probabilities (e.g., in phylogenetic inference, Felsenstein, 1985; Yang and Rannala, 1997; Huelsenbeck and Ronquist, 2001; Heled and Drummond, 2009; Lemoine et al., 2018; Meyer et al., 2019). While Bayesian implementations and other methods to approximate confidence intervals around the predictions exist for DL models, they are not easily scalable for large models or offer limited robustness in the estimation of uncertainties (Blundell et al., 2015; Gal and Ghahramani, 2016; Polson and Sokolov, 2017; Silvestro and Andermann, 2020). Part of the reason for this stems from the fact that artificial intelligence research, unlike evolutionary biology, has for the most part focused on accuracy scores rather than on the estimation uncertainty (Koch et al., 2021).

Third, hypothesis testing is a crucial aspect in evolutionary biology and this is more directly implemented within a probabilistic framework. The statistical comparison between alternative hypotheses typically involves a probabilistic approach, which does not easily have an equivalent in machine learning. Furthermore, delving into the significance of different nodes within a network and comprehending their influence on model performance with a specific dataset assumes an elevated level of complexity. The intricate and nonlinear decision boundaries that are inherent in deep neural networks combined with their extensively parameterized architecture foster an impressive predictive accuracy, but also contribute to the challenge of interpreting them compared to other likelihood-based models.

### Toward a semi-supervised approach to phylogenetic inference

In light of these considerations, we propose that coupling DL with likelihood-based methods can result in more accurate, robust and interpretable estimations of macroevolutionary parameters. We showed that DL can provide accurate estimates of evolutionary rates that are generally considered as nuance parameters in phylogenetic inference (Yang, 1994). Supervised models can efficiently predict these parameters (e.g. per site rates), which can then feed into a likelihood-based inference of the phylogenetic tree. An integrated approach combining DL and likelihood-based inference, can be seen as a form of semi-supervised learning (Zhu, 2005), in which supervised and unsupervised parts of the overall model are applied to different sets of parameters.

The application of this approach to phylogenetic inference reduces the number of free parameters for a likelihood model to optimize, thus reducing its computing costs. For example, gamma or free-rates models typically require re-computing the likelihood of the data at least four times at each optimization or MCMC iteration, i.e. once for each rate category. If per-site rates are instead independently and quickly predicted through a supervised model, the likelihood only needs to be computed one time for each iteration. Our experiments showed that this approach can also potentially improve the accuracy of likelihood-based phylogenetic estimation, by providing higher resolution of rate heterogeneity, and more frequently favoring the true tree against alternative ones. Thus, we envision a new generation of semi-supervised phylogenetic models that integrate likelihood-based and DL components to improve model efficiency and scalability, while relaxing some of the assumptions currently made for mathematical convenience, paving the way for a better understanding of macro-evolutionary processes.

## Acknowledgements and funding

We thank Anna Marcionetti for help with the clownfish data and Diego A. Hartasánchez for feedback on the manuscript.

D.S. received funding from the Swiss National Science Foundation (PCEFP3_187012), the Swedish Research Council (VR: 2019-04739), and the Swedish Foundation for Strategic Environmental Research MISTRA within the framework of the research programme BIOPATH (F 2022/1448).

N.S. and T.L. were supported by a grant from the Swiss National Science Foundation (310030_185223) to N.S. and funding from the University of Lausanne.

## Supplementary Information

### Supplementary tables

**Table S1:** Parameter of the evolutionary models. Note that base frequencies only apply to HKY and GTR models the transition-transversion ratio (Ts / Tv) only applies to the HKY model, while the instantaneous rates only apply to the GTR model.

| Parameter | Distribution |
| --- | --- |
| Substitution model | {JC, HKY, GTR} |
| Base frequencies | $\text{Dir}(\alpha, \alpha, \alpha, \alpha)$, with $\alpha = 5$ |
| Instantaneous rates | $6 \times \text{Dir}(\alpha, \alpha, \alpha, \alpha, \alpha, \alpha)$, with $\alpha = 5$ |
| Ts / Tv ratio | $\mathcal{U}(2, 12)$ |

**Table S2:** Log-likelihood under the true tree. 600 simulations (rows) under different heterogeneity models and tree length. For each simulation, Log-likelihood of the simulated data is computed given the true tree, given the nucleotide matrix estimated under gamma rates, and given site-specific rates. Site-specific rates are either posteriors under gamma model, posteriors under free-rates model, estimated by phyloRNN or finally the true rates (used as input of the simulation). The full table (600 rows) is available at github.com/phyloRNN/SupplementaryMaterials/blob/main/table_S2.csv

| Replicate | Heterogeneity model | Tree length | LnL gamma rates | LnL free-rates | LnL phyloRNN rates | LnL true rates |
| --- | --- | --- | --- | --- | --- | --- |
| 0 | GBM uncorrelated | 0.048 | -1735.990 | -1735.770 | -1717.994 | -1735.399 |
| 1 | Gamma autocorrelated | 0.617 | -5252.660 | -5246.910 | -5055.227 | -5095.449 |
| 2 | Bimodal uncorrelated | 0.235 | -2665.980 | -2664.470 | -2585.512 | -2573.566 |
| 3 | Spike-and-slab uncorrelated | 0.560 | -4718.620 | -4713.730 | -4625.221 | -4686.640 |
| 4 | GBM uncorrelated | 0.064 | -1895.890 | -1877.200 | -1797.892 | -1800.167 |
| 5 | Codon uncorrelated | 1.189 | -7821.070 | -7817.960 | -7488.423 | -7497.536 |
| 6 | GBM uncorrelated | 16.565 | -17096.470 | -16919.200 | -16074.284 | -15844.467 |
| 7 | Bimodal uncorrelated | 4.568 | -17651.340 | -17359.680 | -16690.113 | -16597.409 |
| 8 | Bimodal uncorrelated | 2.719 | -13303.310 | -13201.690 | -12596.996 | -12568.510 |
| 9 | GBM uncorrelated | 3.089 | -14481.340 | -14437.670 | -13647.083 | -13678.580 |
| 10 | Spike-and-slab autocorrelated | 5.565 | -23700.900 | -23694.690 | -23443.449 | -23478.195 |
| ⋮ | ⋮ | ⋮ | ⋮ | ⋮ | ⋮ | ⋮ |

### Supplementary figures

**Figure S1:**
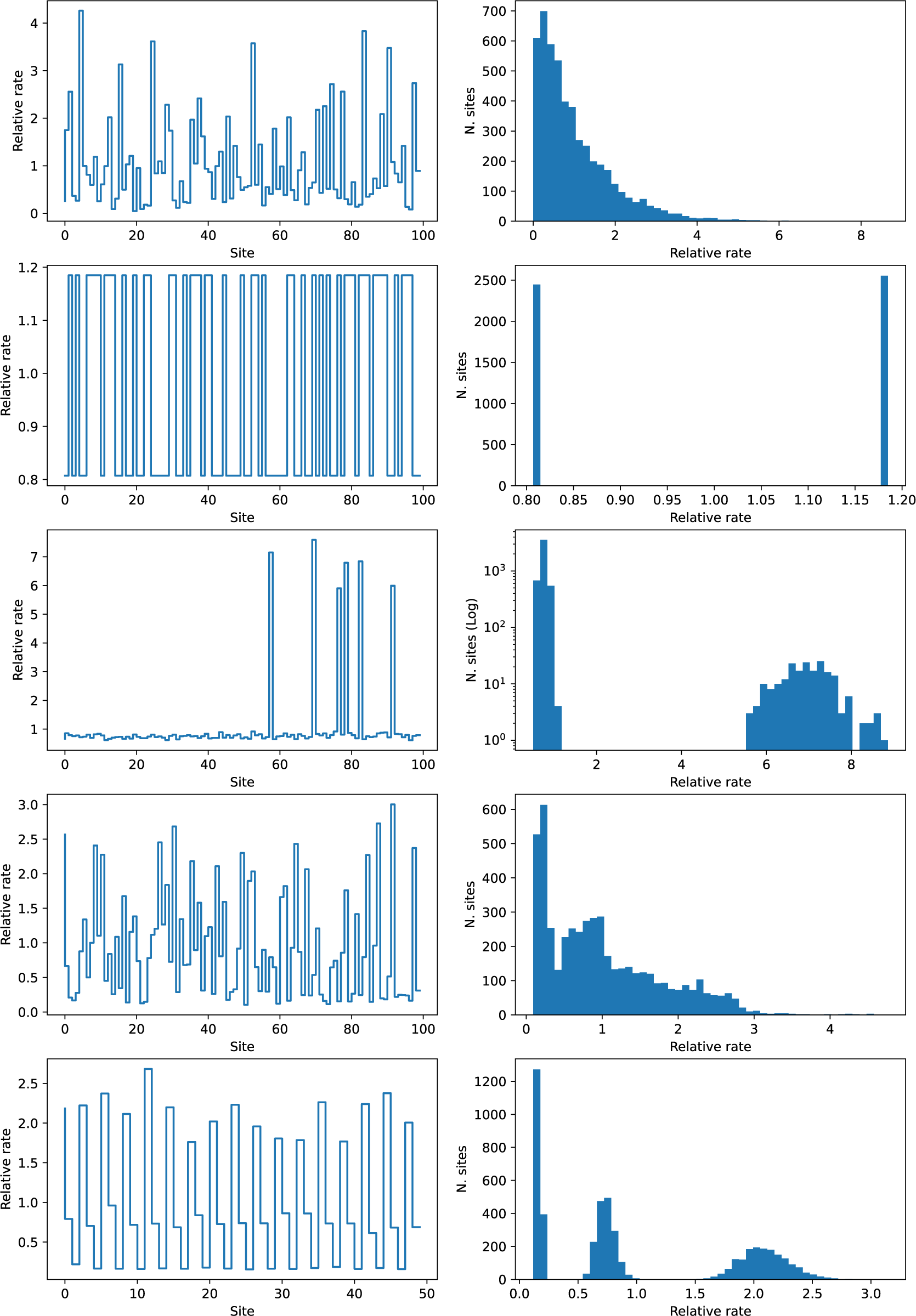
Examples of site-specific rates (plots on the left) and their distribution (histograms on the right) based on simulations with 5,000 sites generated under different modes of rate heterogeneity : a) gamma, b) bimodal, c) spike-and-slab, d) geometric Brownian, e) codon (see Text for more details). Note that for clarity a variable number of sites are shown in the plots and the Y-axis in histogram c) is log-transformed.

**Figure S2:**
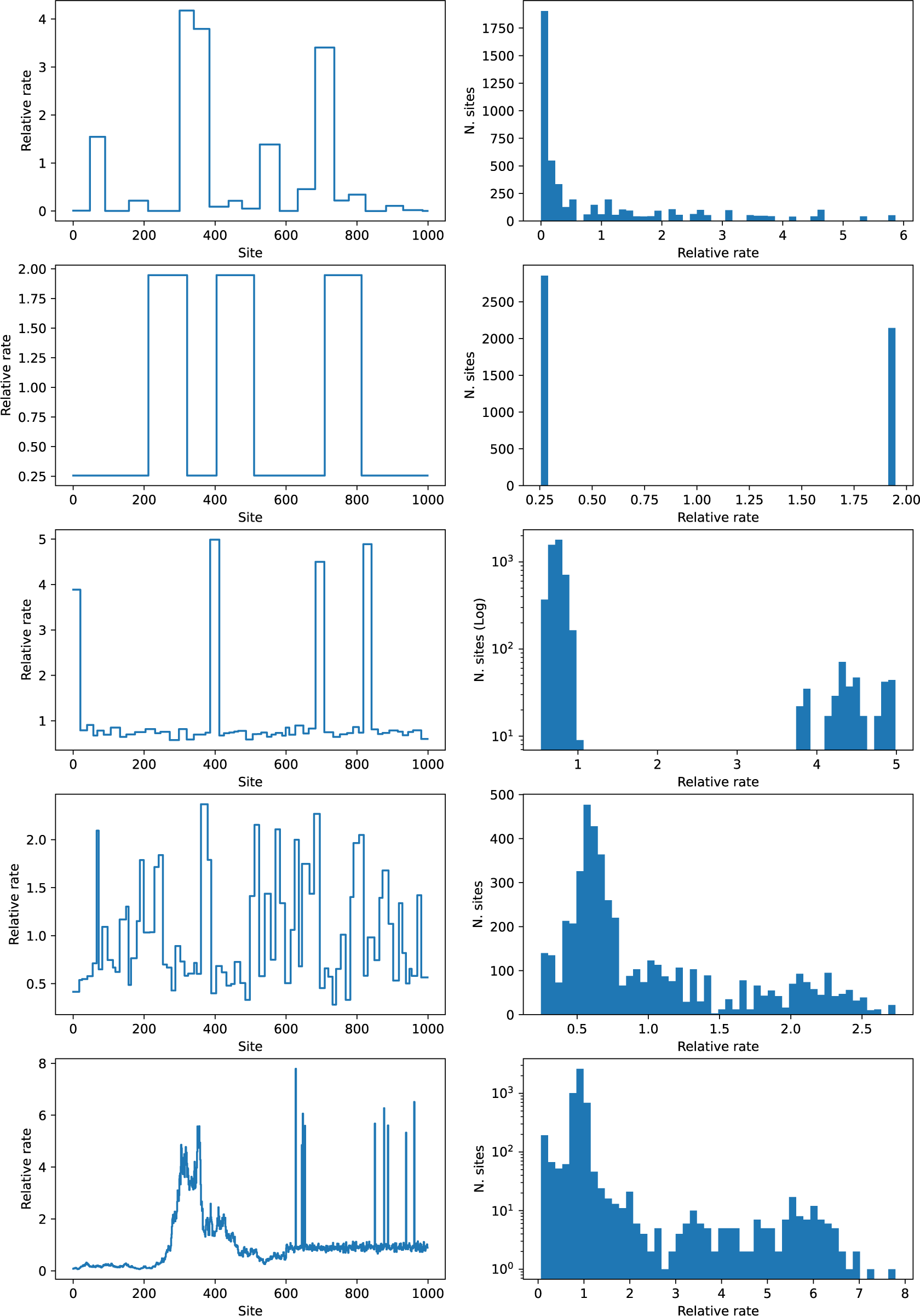
Examples of site-specific rates (plots on the left) and their distribution (histograms on the right) based on simulations with 5,000 sites generated under different autocorrelated modes of rate heterogeneity: a) gamma, b) bimodal, c) spike-and-slab, d) geometric Brownian, e) mixed (see Text for more details). Note that for clarity the Y-axis in histograms c) and e) is log-transformed.

**Figure S3:**
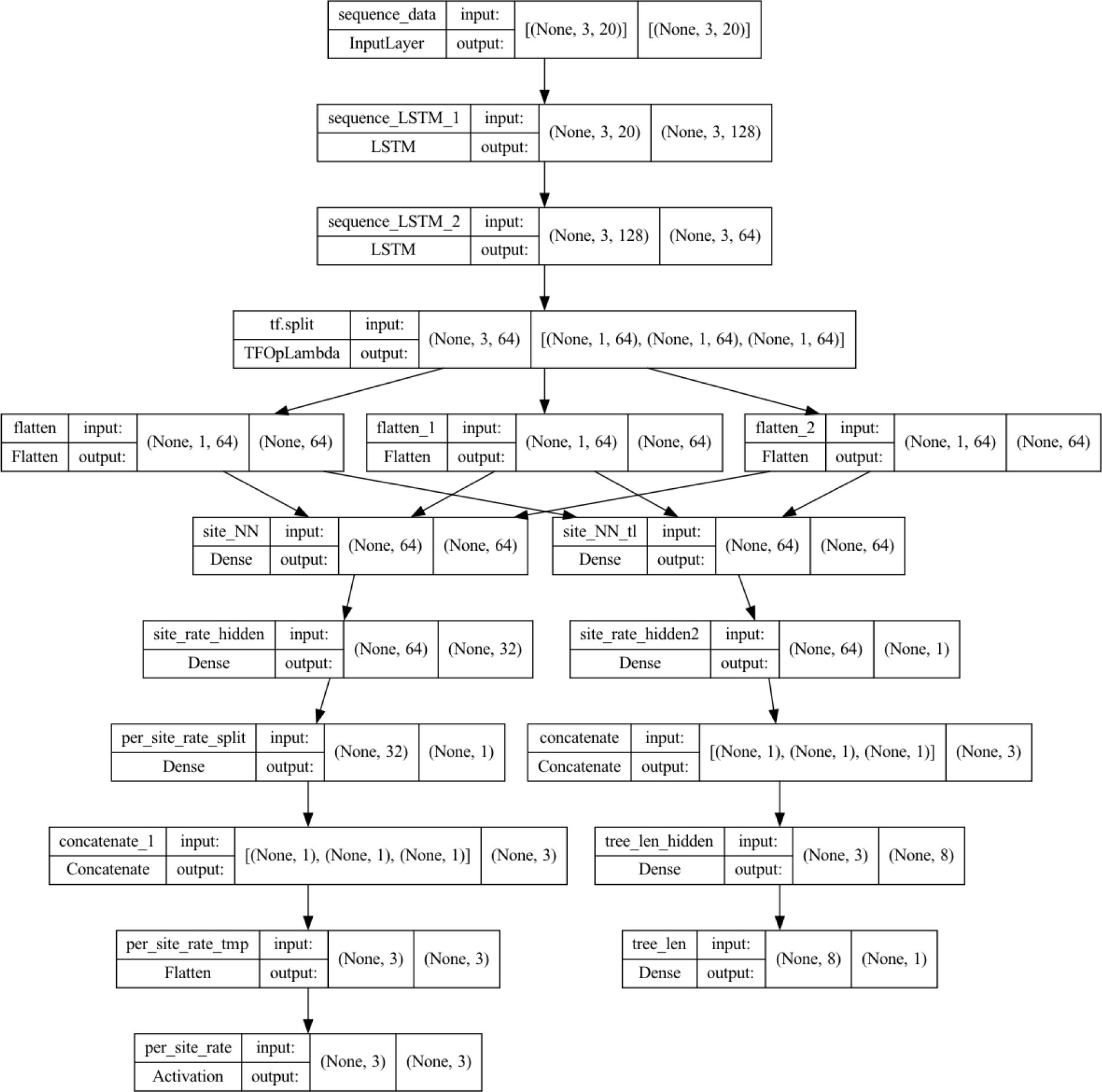
Architecture of the DL model used to infer site-specific rates and total tree length, here show for a toy example with the input being an alignment of 5 sequences of 3 sites. The shape of the input is: (n. sites = 3, n. sequences *×* 4 = 20, with 4 representing the four one-hot-encoded nucleotides. The first dimension indicated with None represents the number of instances, e.g. the number of alignments in the training or test sets.

**Figure S4:**
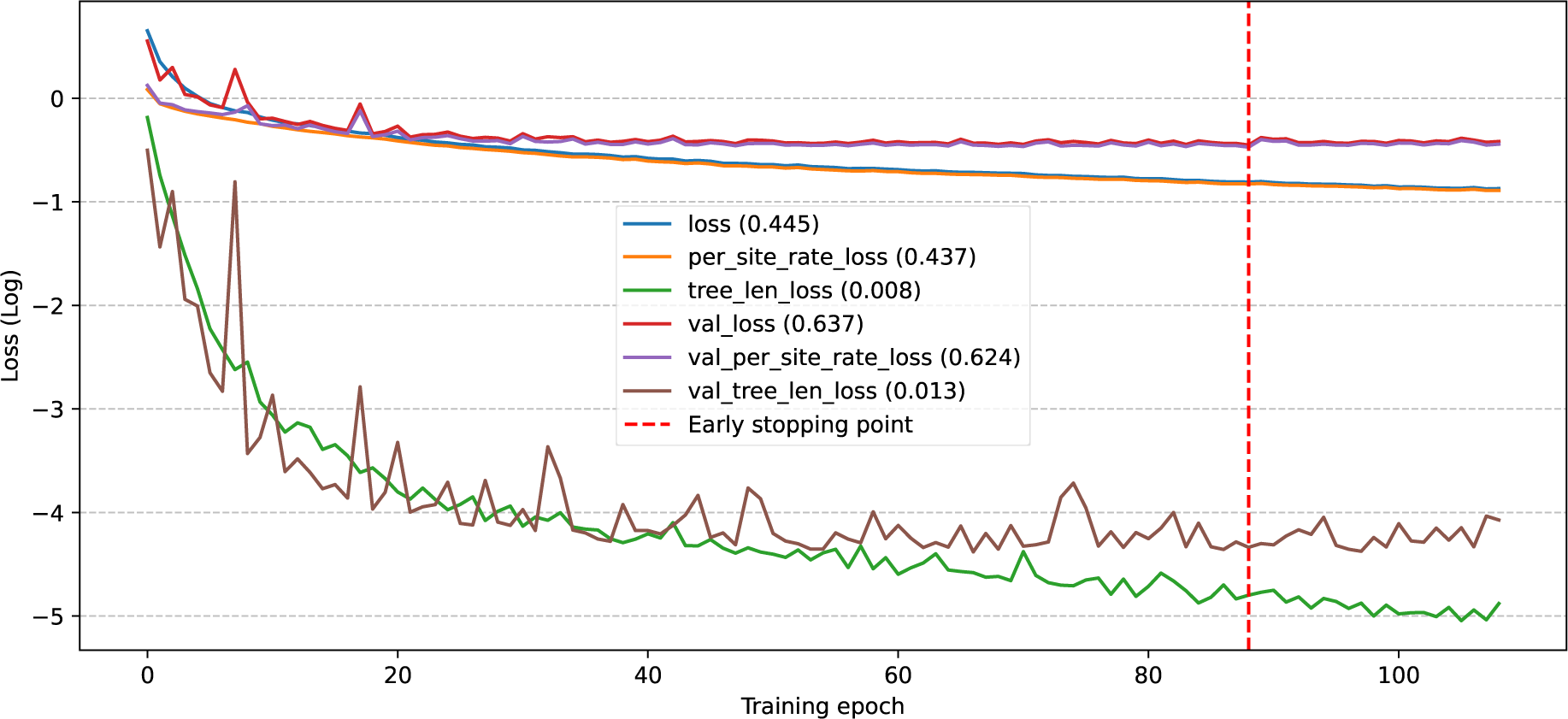
Training history of our DL model. The early stopping point reflects the epoch with the lowest validation loss, which combined the mean squared error of the per-site relative substitution rates and the mean squared error of the total tree length (log-transformed).

**Figure S5:**
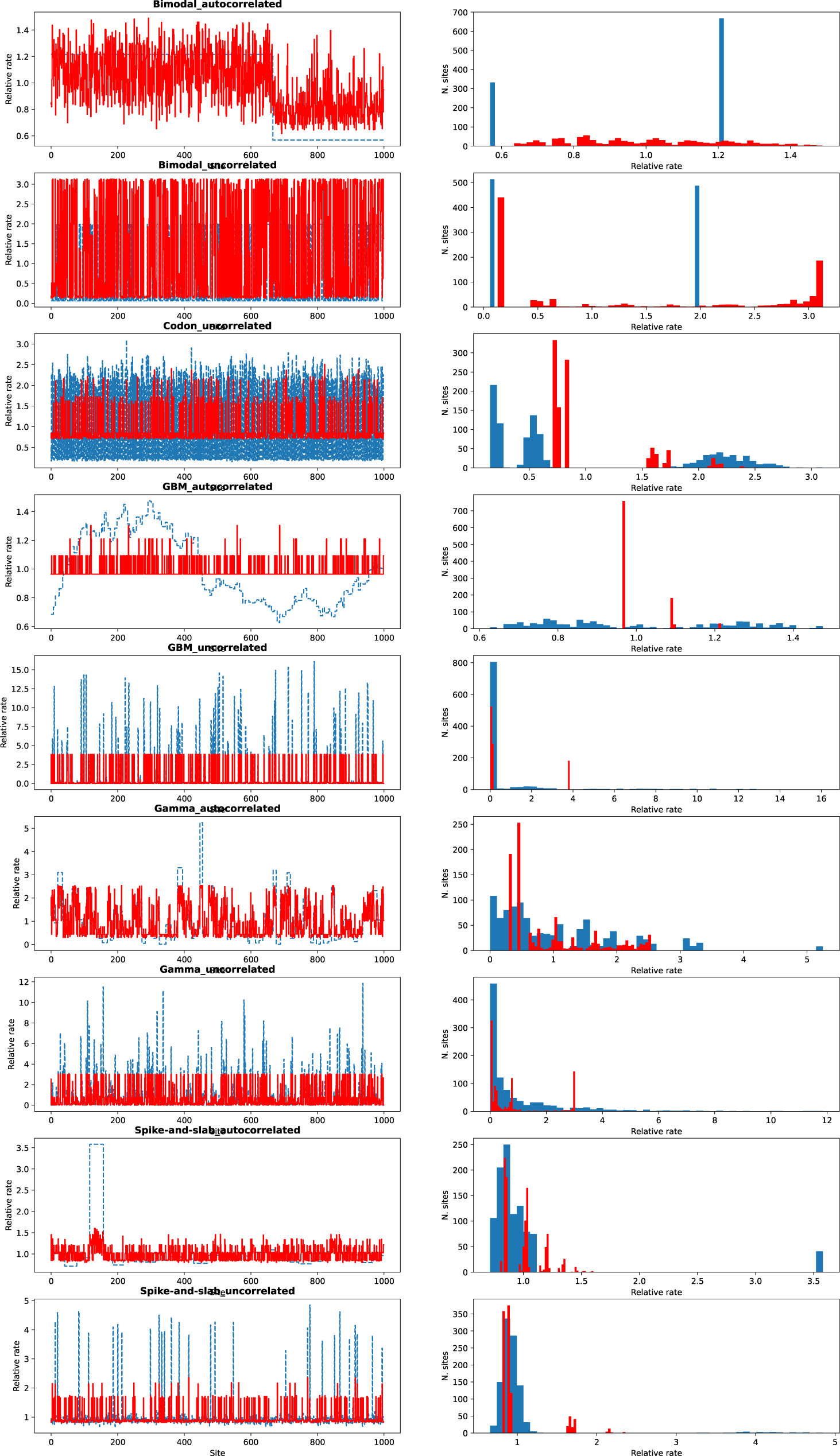
Examples of estimated rates (red) obtained under a maximum likelihood framework using a gamma model of rate heterogeneity. True rates are shown in blue. the estimates are shown as per-site rate (1,000 sites in each dataset) and as histograms.

**Figure S6:**
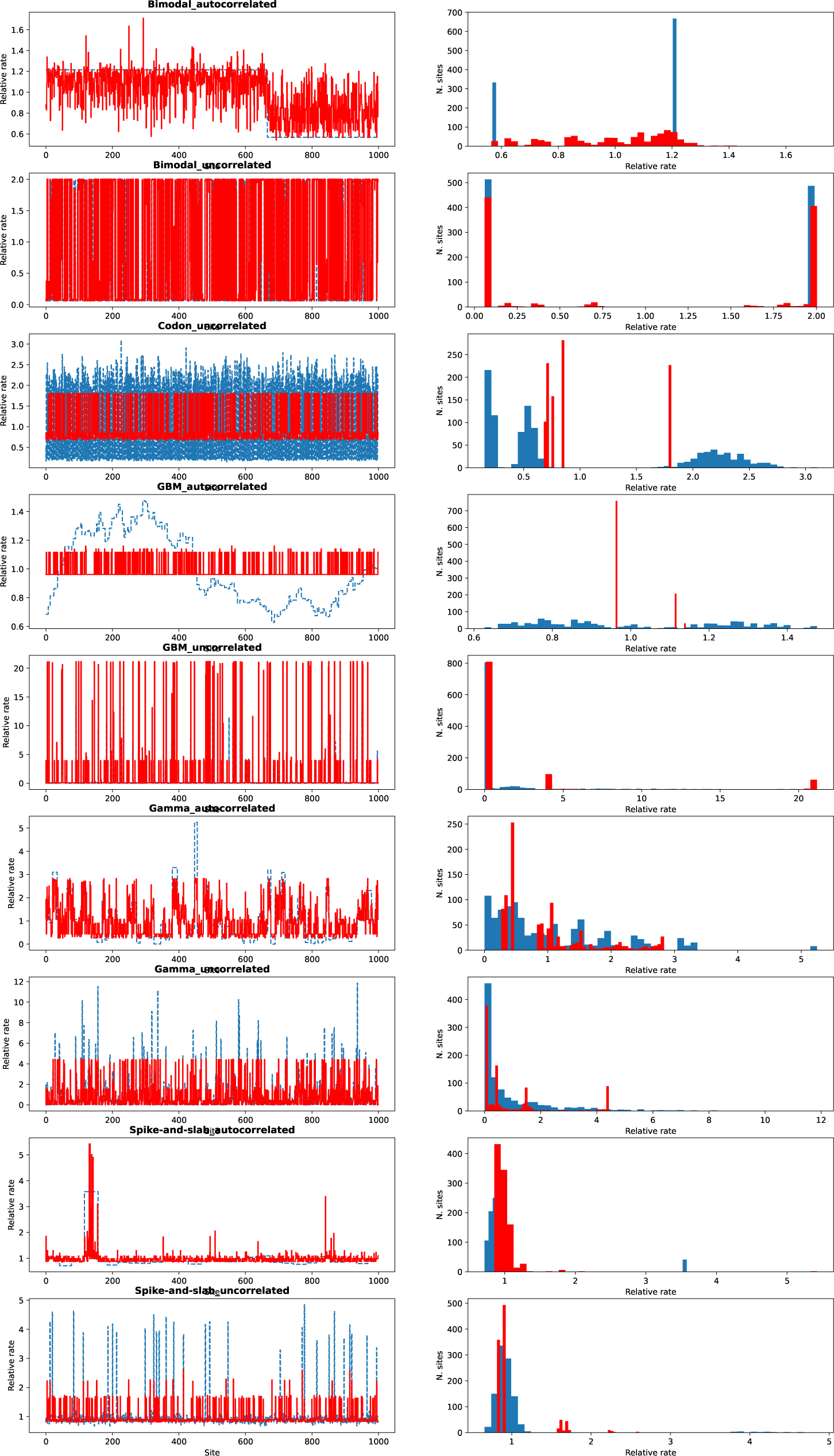
Examples of estimated rates (red) obtained under a maximum likelihood framework using a free-rates model of rate heterogeneity. True rates are shown in blue. the estimates are shown as per-site rate (1,000 sites in each dataset) and as histograms.

**Figure S7:**
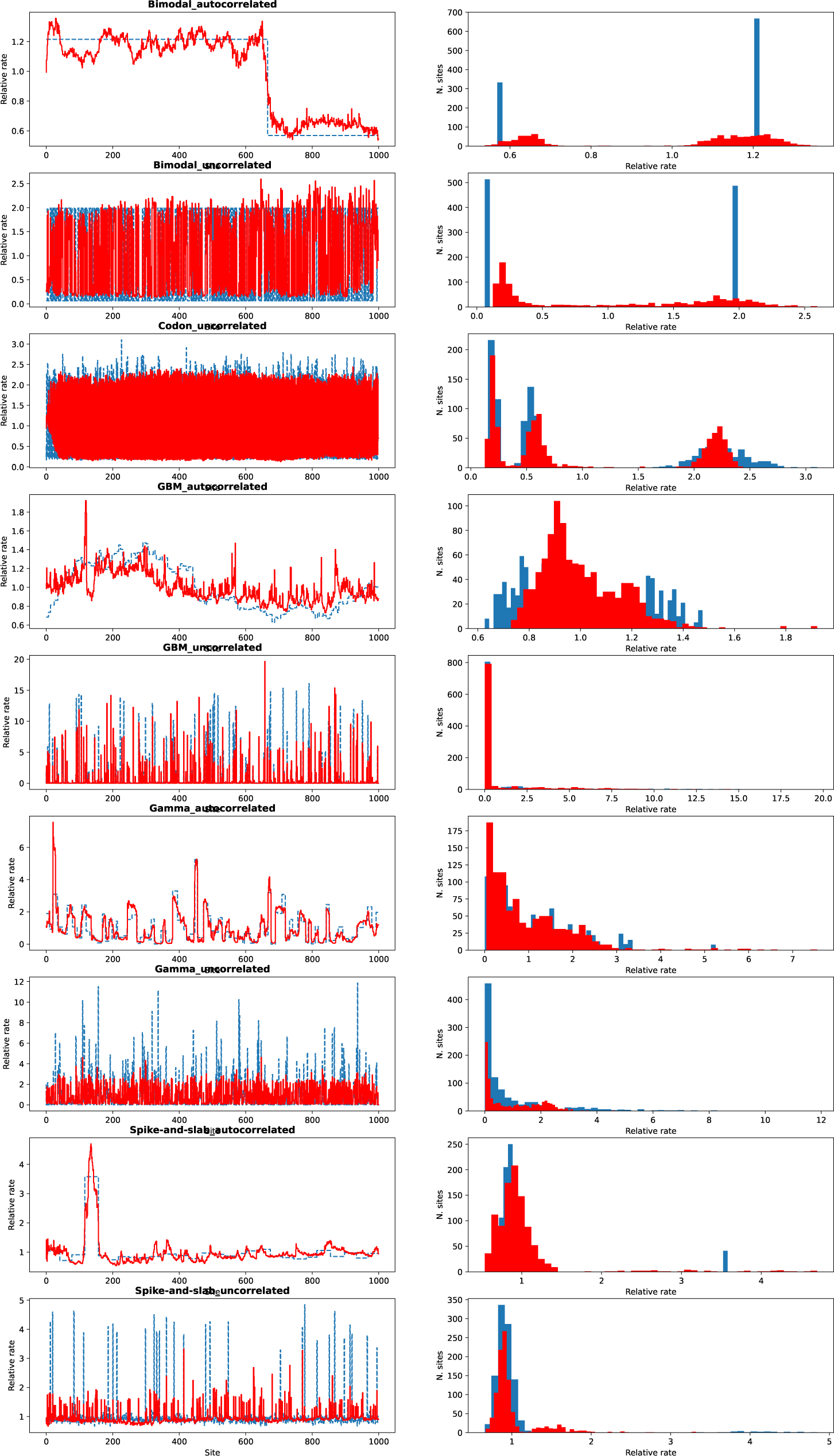
Examples of estimated rates (red) obtained under our DL framework using the phyloRNN model. True rates are shown in blue. the estimates are shown as per-site rate (1,000 sites in each dataset) and as histograms.

**Figure S8:**
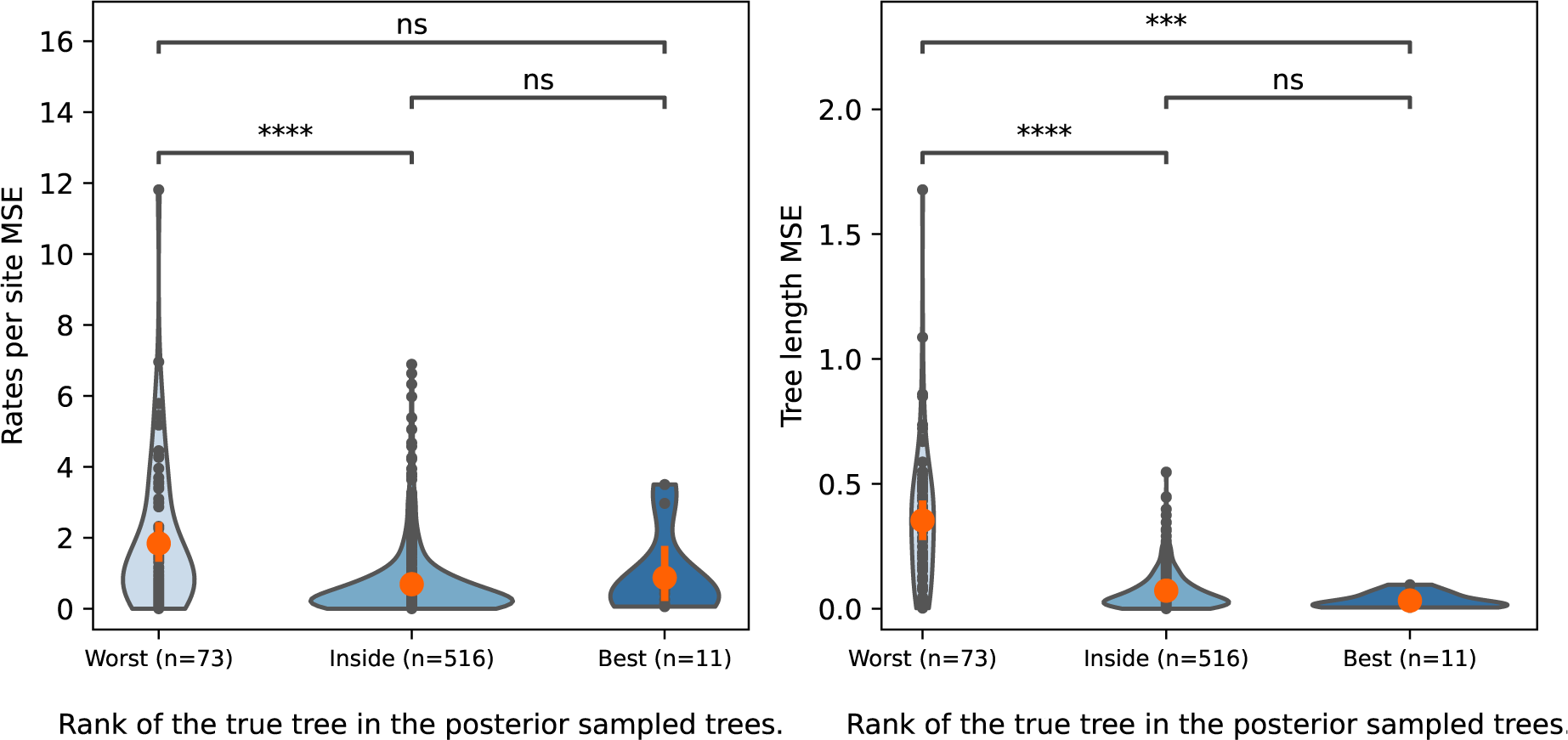
Comparison of Mean-squared error (MSE) across 600 simulations between true values and estimated values (under gamma model). MSE for the rates per site (left) and tree length (right). For each simulation, the likelihood of the true tree is compared to the likelihoods of a posterior sample of 50 trees obtained from a Bayesian analysis (GTR + gamma model). Each of 600 simulations is thus classified whether the true tree is the worst (n=73), inside (n=516) or the best (n=11) of the posterior sampled trees. Comparison of MSE between each pairs of class (worst, inside, best) is performed with t-test of independence. P-values accounting for Bonferroni correction; ns: 0.05 < p <= 1; ***: 1e-04 < p <= 1e-03; ****: p <= 1e-04. Mean of MSE for each class (worst, inside, best) is represented as a red dot.

